# An integrative study on ribonucleoprotein condensates identifies scaffolding RNAs and reveals a new player in Fragile X-associated Tremor/Ataxia Syndrome

**DOI:** 10.1101/298943

**Authors:** Fernando Cid-Samper, Mariona Gelabert-Baldrich, Benjamin Lang, Nieves Lorenzo-Gotor, Riccardo Delli Ponti, Lies-Anne WFM Severijnen, Benedetta Bolognesi, Ellen Gelpi, Renate K. Hukema, Teresa Botta-Orfila, Gian Gaetano Tartaglia

**Affiliations:** Centre for Genomic Regulation (CRG), The Barcelona Institute for Science and Technology, Dr. Aiguader 88, 08003 Barcelona, Spain; Department of Clinical Genetics, Erasmus MC, 3000 CA Rotterdam, The Netherlands; Neurological Tissue Biobank of the Hospital Clinic and Institut d’Investigacions Biomèdiques August Pi i Sunyer (IDIBAPS), Carrer del Rosselló, 149, 08036, Barcelona, Spain; Institute of Neurology, Medical University of Vienna, Währinger Gürtel 18-20 1090 Vienna, Austria; DNA and Fluids Biobank of the Institut d’Investigacions Biomèdiques August Pi i Sunyer (IDIBAPS), Carrer Rosselló 149, 08036 Barcelona, Spain; Universitat Pompeu Fabra (UPF), 08003 Barcelona, Spain; Department of Biology ‘Charles Darwin’, Sapienza University of Rome, P.le A. Moro 5, Rome 00185, Italy; Institució Catalana de Recerca i Estudis Avançats (ICREA), 23 Passeig Lluís Companys, 08010 Barcelona, Spain

**Keywords:** Phase separation, Scaffolding RNA, CGG repeat expansion, FMR1 premutation, Fragile X-associated tremor/ataxia syndrome (FXTAS), RNA aggregates, RNA binding proteins (RBP), TRA2A splicing regulator, Neurodegeneration

## Abstract

Recent evidence indicates that specific RNAs promote formation of ribonucleoprotein condensates by acting as scaffolds for RNA-binding proteins (RBPs).

We systematically investigated RNA-RBP interaction networks to understand ribonucleoprotein assembly. We found that highly-contacted RNAs are highly structured, have long untranslated regions (UTRs) and contain nucleotide repeat expansions. Among the RNAs with such properties, we identified the *FMR1* 3’ UTR that harbors CGG expansions implicated in Fragile X-associated Tremor/Ataxia Syndrome (FXTAS).

We studied *FMR1* binding partners *in silico* and *in vitro* and prioritized the splicing regulator TRA2A for further characterization. In a FXTAS cellular model we validated TRA2A-*FRM1* interaction and investigated implications of its sequestration at both transcriptomic and post-transcriptomic levels. We found that TRA2A co-aggregates with *FMR1* in a FXTAS mouse model and in *post mortem* human samples.

Our integrative study identifies key components of ribonucleoprotein aggregates, providing links to neurodegenerative disease and allowing the discovery of new therapeutic targets.

## Introduction

Proteins and RNAs coalesce in large phase-separated condensates that are implicated in several cellular processes (Jiang et al., 2015; Woodruff et al., 2017).

Among the most studied condensates, ribonucleoprotein (RNP) granules assemble in liquid-like cellular compartments composed of RNA-binding proteins (RBPs) (Hyman et al., 2014; Maharana et al., 2018) that are in dynamic exchange with the surrounding environment (Bolognesi et al., 2016). RNP granules such as processing bodies and stress granules (SGs) are evolutionarily conserved from yeast to human (Brangwynne et al., 2009; Jain et al., 2016; Riback et al., 2017) and contain constitutive protein components such as G3BP1 (yeast: Nxt3), TIA1 (Pub1), and TIAR (Ngr1) (Buchan et al., 2008). Several granule-forming RBPs are prone to form amyloid aggregates upon amino acid mutations (Hyman et al., 2014; Kato et al., 2012) that induce a transition from a liquid droplet to a solid phase (Qamar et al., 2018). This observation has led to the proposal that a liquid-to-solid phase transition is a mechanism of cellular toxicity (Patel et al., 2015) in diseases such as Amyotrophic Lateral Sclerosis (ALS) (Murakami et al., 2015) and Myotonic Dystrophy (Pettersson et al., 2015).

All the components of molecular complexes need to be physically close to each other to perform their functions. One way to achieve this, while keeping selectivity in a crowded cell, is to employ platform or scaffold molecules that piece together components of a complex or a pathway. Indeed, RBPs are known to act as scaffolding elements promoting RNP assembly through protein-protein interactions (PPI) (Banani et al., 2017), yet protein-RNA interactions (PRI) also play a role in the formation of condensates. Indeed, a recent work based on G3BP1 pull-down indicates that 10% of the human transcripts can assemble into SGs (Khong et al., 2017). If distinct RNA species are present in the condensates, a fraction of them could be involved in mediating RBP recruitment. In this regard, we previously observed that different RNAs act as scaffolds for RNP complexes (Ribeiro et al., 2018), which indicates that specific transcripts might promote formation of RNP condensates.

Combining PPI and PRI networks revealed by enhanced CrossLinking and ImmunoPrecipitation (eCLIP) (Van Nostrand et al., 2016a) and mass spectrometric analysis of SGs (Jain et al., 2016), we identified a class of transcripts that bind to a large number of proteins and, therefore, qualify as potential scaffolding elements. In agreement with recent literature reports, we found that untranslated regions (UTRs) have a particularly strong potential to bind proteins in RNP granules, especially when they contain specific homo-nucleotide repeats (Saha and Hyman, 2017). In support of this observation, several diseases including Myotonic Dystrophy (DM) and a number of ataxias (SCA) have been reported to be linked to expanded trinucleotide repeats that trigger formation of intranuclear condensates in which proteins are sequestered and functionally impaired. Specifically, expanded RNA repeats lead to RNA-mediated condensate formation in DM1 (Mooers et al., 2005), SCA8 (Mutsuddi et al., 2004) and SCA10 (White et al., 2010).

By understanding the characteristics of RNAs involved in RNP assembly we aim to unveil the molecular details of specific human diseases. Indeed, appearance of RNP condensates, often called *inclusions* or *foci*, is not only linked to ALS, Huntington’s disease and Myotonic Dystrophy, but also other diseases such as Fragile X-associated Tremor / Ataxia Syndrome (FXTAS) (Tassone et al., 2004; Sellier et al., 2017). The onset and development of FXTAS is currently explained by two main mechanisms (Botta-Orfila et al., 2016): i) RNA-mediated recruitment of proteins attracted by CGG trinucleotide repeats in the 5’ UTR of Fragile X Mental Retardation Protein (*FMR1*) RNA and ii) aggregation of Repeat-Associated Non-AUG (RAN) polyglycines peptides translated from the *FMR1* 5’ UTR (FMRpolyG) (Todd et al., 2013). Previous work indicates that *FMR1* inclusions contain specific proteins such as HNRNP A2/B1, MBNL1, LMNA and INA (Iwahashi et al., 2006). Also FMRpolyG peptides (Sellier et al., 2017) have been found in the inclusions, together with CUGBP1, KHDRBS1 and DGCR8 that are involved in splicing regulation, mRNA transport regulation of microRNA regulation (Sellier et al., 2010, 2013). While KHDRBS1 does not bind physically (Sellier et al., 2010), its protein partner DGCR8 interacts with CGG repeats (Sellier et al., 2013), indicating that sequestration is a process led by a pool of proteins that progressively attract other networks.

Notably, CGG repeats contained in the FMR1 5’ UTR are of different length (the most common allele in Europe being of 30 repeats). At over 200 repeats, methylation and silencing of the *FMR1* gene block FMRP protein expression (Todd et al., 2013). The premutation range (55–200 CGG repeats) is instead accompanied by appearance of *foci* that are the typical hallmark of FXTAS (Todd et al., 2013). These *foci* are highly dynamic and behave as RNP condensates that phase separate in the nucleus forming inclusions (Tassone et al., 2004). Although long-lived, they rapidly dissolve upon tautomycin treatment, which indicates liquid-like behavior (Strack et al., 2013).

The lability of *FMR1* inclusions, which impedes their biochemical characterization (Mitchell et al., 2013; Marchese et al., 2016), complicates the identification of RBPs involved in FXTAS. As shown in previous studies of ribonucleoprotein networks (Cirillo et al., 2017; Marchese et al., 2017), computational methods can be exploited to identify key partners of RNA molecules. New contributions from other research areas are needed, especially because FXTAS pathological substrate is still under debate and there is still insufficient knowledge of targets for therapeutic intervention (Todd et al., 2013; Sellier et al., 2017). We here propose an integrative approach to identify new markers based on properties of PRI networks and characteristics of scaffolding RNAs.

## Results

In this work we exploited a high-throughput computational approach to investigate the physico-chemical properties of scaffolding RNAs (Figure 1A). We focused on the experimental characterization of *FMR1* that we predict to bind a large number of RBPs. Among the *FMR1* partners that we identified, we selected the splicing regulator TRA2A and studied the biological consequences of its recruitment in RNP condensates. We used murine and *post mortem* human tissues to assess TRA2A involvement in FXTAS.

**Figure 1.**
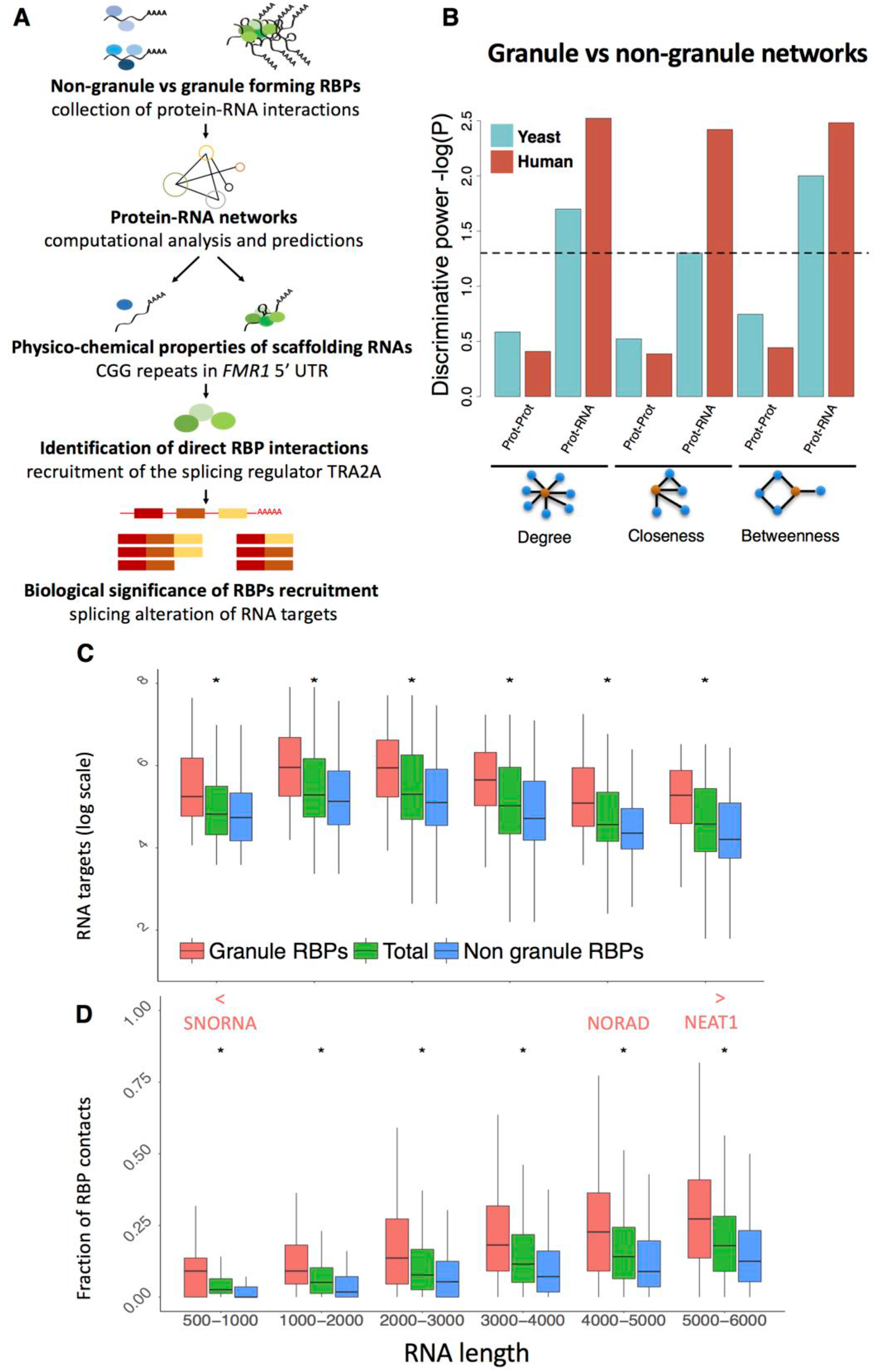
RNA as a key element in RNP condensates. **A)** We explored the differences between granule and non-granule RBPs using an interaction network approach. We first studied the physico-chemical properties of scaffolding RNAs and prioritized *FMR1* 5’ UTR for experimental characterization. We retrieved known RBPs and identified *FMR1* partners involved in FXTAS, including splicing factors. We evaluated the biological consequences of RBP recruitment in cellular context and the presence of RNP complexes in FXTAS brain inclusions. **B)** Statistical differences between granule and non-granule elements (proteins or RNA) in PPI and PRI networks. Only when analyzing RNA interactions, granule and non-granule networks show different topologies (**Supplementary Figure 1C, 1D**). **C)** Independently of RNA length, granule RBPs contact more transcripts that non-granule RBPs (‘total’ indicates all RBPs regardless of their granule or non-granule definition; * indicates p-value < 0.01; Kolmogorv-Smirnov, KS, test). **D)** Transcripts interact more frequently with granule than non-granule RBPs. The fractions of granule and non-granule RBP contacts, monitored at different lengths, shows consistent enrichments (p-value < 0.01; KS test). Highly contacted transcripts are enriched in small nuclear (snRNAs) and small nucleolar RNAs (snoRNAs) (p-value < 2.2e-16, Wilcoxon test). Already described scaffolding RNAs such as NEAT1 are also identified.

### Protein-protein networks do not discriminate granule and non-granule RBPs

We first studied if RBPs phase-separating in RNP condensates interact with specific sets of proteins and RNAs. To discriminate proteins that are in RNP condensates (granule RBPs) from other RBPs (non-granule RBPs) we relied on recent proteomics data on human and yeast SGs (**STAR Methods** and **Table S1A**) as well as computational methods. The protein-RNA interaction datasets were identified through eCLIP (human) (Van Nostrand et al., 2016a) and microarray (yeast) (Mittal et al., 2011) studies (**Table S1B**).

We analyzed if granule and non-granule RBPs show different interaction network properties. To this aim, we used available PPI datasets **(STAR Methods**) (Huttlin et al., 2015a; Mittal et al., 2011). We based the topological analysis on three centrality measures describing the importance of a node (protein) within the network. For each protein, we computed the degree (number of protein interactions), betweenness (number of paths between protein pairs) and closeness centrality (how close one protein is to other proteins in the network). We found that granule and non-granule RBPs networks display very similar topology both in yeast and human datasets (Figure 1B; **Supplementary Figure 1A, 1B**).

### Protein-RNA networks robustly discriminate granule and non-granule RBPs

In both yeast and human, we found that PRIs increase significantly the centrality measures of the granule network (Figure 1B and **Supplementary Figure 1C, 1D**). Importantly, human and yeast granule RBPs interact with more transcripts than other RBPs (Figure 1C; **Supplementary Figure 2; Tables S1C, S1D, S1E** and **S1F**; p-value yeast = 0.02, p-value human = 0.003, Wilcoxon test; **STAR Methods**). Such a difference holds even when looking independently at either coding or non-coding RNAs (**Supplementary Figure 2;** p-value coding = 0.003, p-value non-coding = 0.01, Wilcoxon test) and upon normalization by transcript length (p-value yeast = 0.02; p-value human = 0.002, Wilcoxon test).

### Granule RBPs share RNA networks

In both yeast and human proteomes we found that granule-forming RBPs share a larger number of transcripts (**Supplementary Figure 3A; Tables S1G, S1H, S1I** and **S1J;** p-value yeast < 2.2e-16, p-value K562 < 2.2e-16, KS test). Independently of their length, RNAs contacting granule RBPs preferentially interact with other granule RBPs (Figure 1D, p-value < 2.2e-16, Wilcoxon test). In agreement with this finding, RNAs interacting exclusively with granule RBPs have a higher number of protein contacts than RNAs associating only with non-granule RBPs (**Supplementary Figure 3B**, p-value = 0.04, Wilcoxon test). This observation is consistent with a picture in which RNAs share a high number of RBPs interactions to promote recruitment into RNP granules. Using a high confidence threshold to select RBP partners (number of reads normalized by expression levels in the third quartile of the statistical distribution) (Armaos et al., 2017a), we found that our list of RNAs overlaps with a recently published atlas of transcripts enriched in SGs (Area under the ROC curve or AUC of 0.89; sensibility of 81.3% and specificity of 85.2%; **Supplementary Figure 3C**; **Table S1K**) (Khong et al., 2017).

### Non-coding RNAs are contacted by granule RBPs

Among the most contacted RNAs we found an enrichment of small nuclear and nucleolar RNAs that are known to be associated with paraspeckles and Cajal bodies formation (Figure 1D; **Table S2A;** p-value < 2.2e-16, Wilcoxon test). We also identified a few highly contacted long non-coding RNAs such as *NEAT1* that interacts with all the proteins present in our dataset (Figure 1D). In agreement with this finding, *NEAT1* has been described as an architectural RNA (West et al., 2016) implicated in scaffolding RBPs (Maharana et al., 2018) for paraspeckle nucleation. We hypothesize that other highly contacted long non-coding RNAs may have similar functions within cytoplasmic RNP granules. For instance, *NORAD*, a recently described long non-coding RNA involved in genomic stability, interacts with the large majority of proteins in our dataset (Lee et al., 2016). *NORAD* has repetitive sequence regions, is activated upon stress, has ability to recruit proteins (Tichon et al., 2016) and aggregates in stress granules (Khong et al., 2017).

### Characteristic features of candidate scaffolding RNAs

We next studied which properties support the scaffolding activity of RNAs within granules. In this analysis, we define as granule transcripts those contacted by a larger number of granule-forming RBPs than non-granule forming RBPs (*viceversa* for non-granule transcripts; **STAR Methods; Table S2B** and **S2C**), we found that RNAs enriched in granule RBP contacts are more expressed (Figure 2A; p-value = 5e-11, Kolmogorv-Smirnov, KS test), structured in UTRs (Figures 2B, and 2C Parallel Analysis of RNA Structure PARS data; p-values = 0.005 and 0.05; we note that the signal is enriched at the 3’ UTRs with p-value < 0.001; the 5’ UTRs is associated with a p-value of 0.02; KS test; **STAR Methods**) and with longer UTRs (Figure 2D; 5’UTR is shown; p-value = 0.005, KS test; 3’ UTR is reported in **Supplementary Figure 3D**). This result, also valid in yeast (**Supplementary Figures 3E** and **3F; Tables 2B and 2C**), is consistent with previous observations that length (Zhang et al., 2015), structure (Reineke et al., 2015) and abundance (Jain and Vale, 2017) contribute to RNA assembly into RNP granules (Khong et al., 2017).

**Figure 2.**
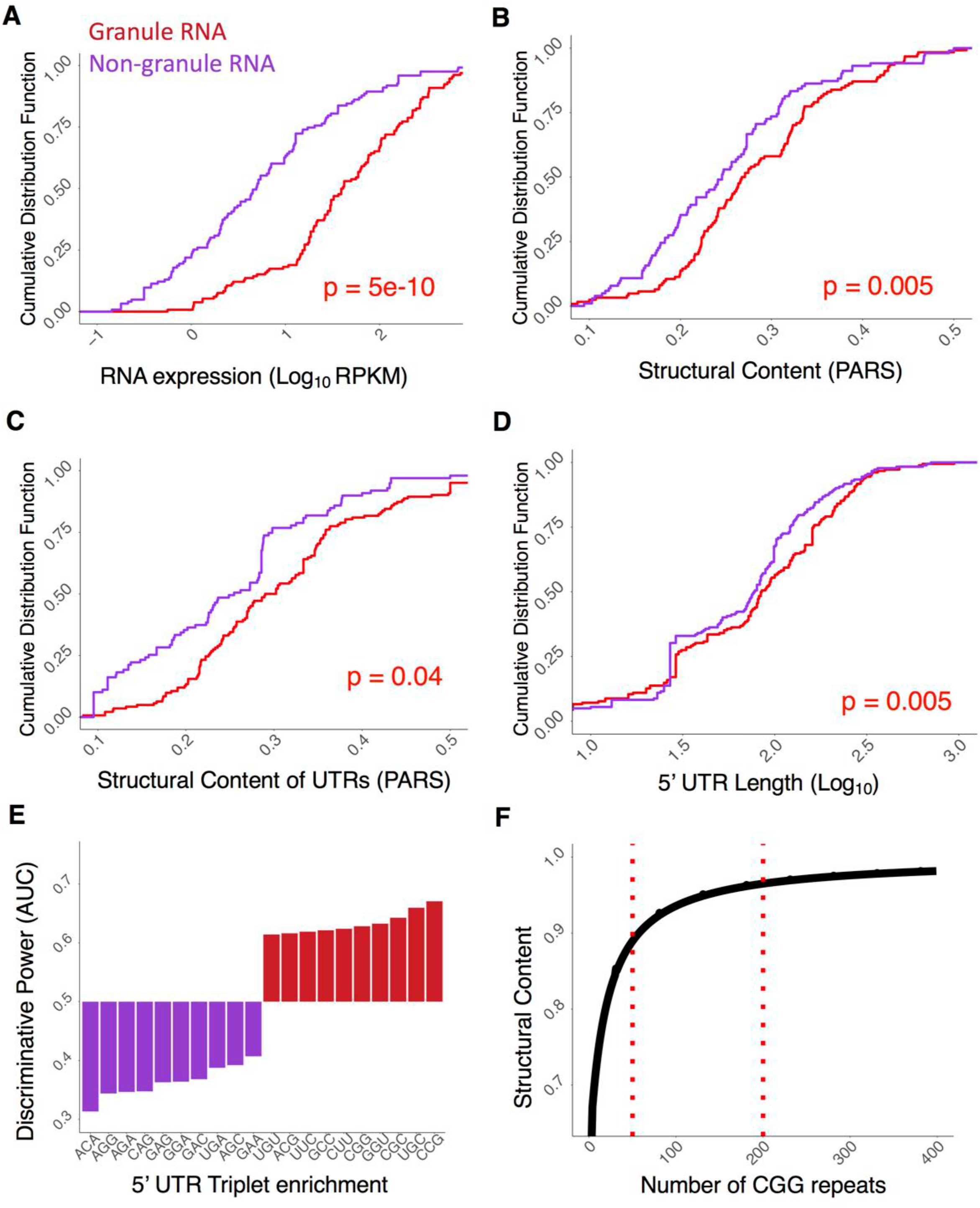
Properties of scaffolding RNAs. **A), B), C)** and **D)** Properties of RNAs contacted by granule-proteins. Granule transcripts are more abundant (**A**, p-value = 4.65e-11, KS test), structured (**B** and **C**, p-value = 0.005 and p-value=0.04; KS test) with longer UTRs (**D**, p-value 5’UTR = 0.005, KS test) than non-granule RNAs. **E)** Occurrence of CCG, UGC, CGC, GGU and CGG repeats discriminate the 5’ UTRs of granule and non-granule transcripts (the Area under the ROC Curve AUC is used to separate the two groups). **F)** Increasing the length of CGG repeats results in stronger secondary structural content (the CROSS algorithm (Delli Ponti et al., 2017) is employed to measure the amount of double-stranded RNA).

Triplets prone to assemble into hairpin-like structures (Krzyzosiak et al., 2012), including CCG, UGC, CGC, GGU and CGG, discriminate granule and non-granule transcripts in the 5’ UTRs (AUCs > 0.60; Figure 2E). In agreement with these findings, predictions of RNA structure performed with the CROSS algorithm (Delli Ponti et al., 2017) indicate that the structural content (presence of double-stranded regions) is enriched in granule-associated transcripts (**Supplementary Figure 4A**) and increase proportionally to CGG repeats length (Figure 2F), which is in line with UV-monitored structure melting experiments (Krzyzosiak et al., 2012).

### In silico predictions indicate a large number of partners for FMR1 scaffolding RNA

To further investigate the scaffolding ability of homo-nucleotide expansions, we selected the *FMR1* transcript that contains CGG repetitions. Using *cat*RAPID *omics* (**STAR Methods**) (Agostini et al., 2013), we computed interactions between the 5’ *FMR1* UTR (containing 79 CGG repeats) and a library of nucleic-acid binding proteins (3340 DNA-binding, RNA-binding and structurally disordered proteins) (Livi et al., 2015). Previously identified CGG-binding proteins (Sellier et al., 2010) such as HNRNP A1, A2/B1, A3, C, D and M, SRSF 1, 4, 5, 6, 7 and 10 as well as MBNL1 and KHDRBS3 were predicted to interact strongly (discriminative power > 0.90) and specifically (interaction strength > 0.90; Figure 3A; **Table S3;** empirical p-values < 0.01). Similar binding propensities were also found for a set of 92 RBPs reported to assemble in SGs (Jain et al., 2016) (**Table S3**). In addition, our calculations identify a group of 37 RBPs that are predicted to form granules by the *cat*GRANULE algorithm (Bolognesi et al., 2016) (**STAR Methods; Supplementary Figures 4B** and **4C**). Among the RBPs prone to associate with *FMR1*, we found a class of splicing factors, including TRA2A (interaction score: 0.99; specificity 1.00; **Table S3**).

**Figure 3.**
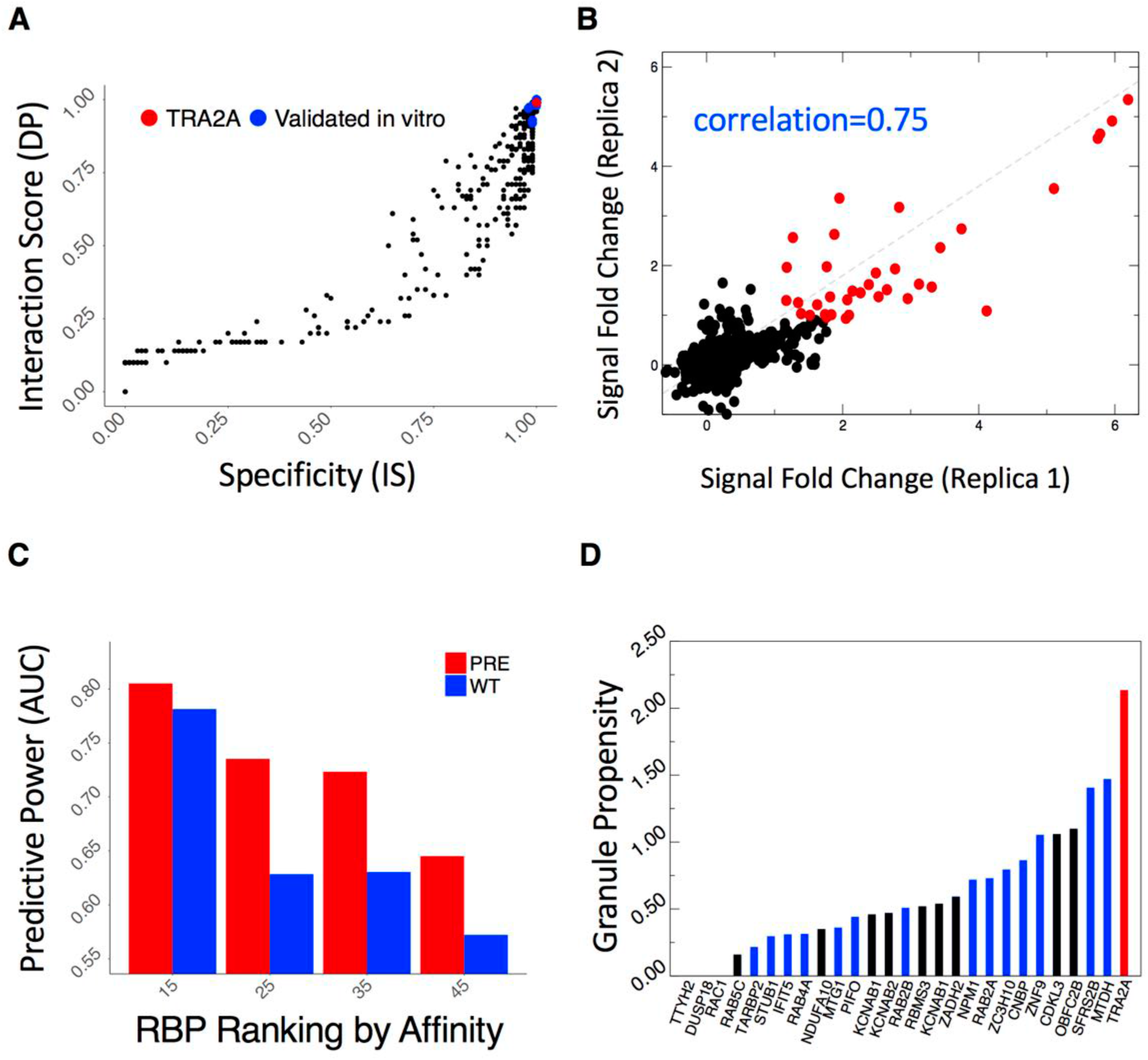
Protein interactions of CGG repeats. **A)** Using *cat*RAPID *omics* (Agostini et al., 2013), we computed protein interactions with the first *FMR1* exon (containing 79 CGG repeats). Previously identified partners, such as HNRNP A1, A2/B1, A3, C, D and M, SRSF 1, 4, 5, 6, 7 and 10 as well as MML1 and KHDRBS3 show strong binding propensities and specificities (blue dots) (Sellier et al., 2010). A previously unknown interactor, TRA2A (red dot) shows comparable binding propensities. **B)** We validated RBP interactions with FMR1 exon (“pre”containing 79 CGG repeats) through protein arrays (Cirillo et al., 2017; Marchese et al., 2017). We obtained high reproducibility between replicas (Pearson’s correlations > 0.75 in log scale) and identified strong-affinity interactions (signal to background ratio > 2.5; red dots). The same procedure was applied to FMR1 exon containing 21 CGG repeats (**Table S4**). **C)** We measured *cat*RAPID *omics* (Agostini et al., 2013) performances on protein array data selecting an equal number of strong-(highest signal to background ratios) and poor-affinity (lowest signal to background ratios) candidates. **D)** Out of 27 candidates binding to both 79 and 21 CGG repeats (signal to background ratio > 2.5), 15 are highly prone to form granules (blue bars) (Bolognesi et al., 2016) and the splicing regulator TRA2A (red bar) shows the highest propensity. The black bars indicate non-specific partners interacting also with SNCA 3’ UTR (Cirillo et al., 2017; Marchese et al., 2017) or showing poor RNA-binding propensity (Livi et al., 2015).

### High-throughput validation of CGG partners and identification of TRA2A interaction

We employed protein arrays (Cirillo et al., 2017; Marchese et al., 2017) to perform a large *in vitro* screening of RBP interactions with the first *FMR1* exon (**STAR Methods**). We probed both expanded (79 CGG) and normal (21 CGG) range repeats on independent replicas, obtaining highly reproducible results (Pearson’s correlations >0.75 in log scale; Figure 3B; **Table S4**). We used the 3’ UTR of a similar length transcript, *SNCA* (575 nt), to control for the specificity of RBP interactions (Marchese et al., 2017).

Using fluorescence intensities (signal to background ratio) to measure binding affinities, we found that previously identified partners SRSF 1, 5 and 6 rank in the top 1% of all interactions (out of 8900 proteins), followed by KHDRBS3 (2%) and MBNL1 (5%). We observed strong intensities (signal to background ratio > 1.5 corresponding to top 1% of all interactions) for 85 RBPs interacting with expanded repeats (60 RBPs for normal-range repeats) and using more stringent cut-offs (signal to background ratio > 2.5 or top 1 ‰ of all interactions) we identified 27 previously unreported interactions (binding to both expanded and normal range repeats).

The list of 85 RBPs showed enrichment in GO terms related to splicing activity (FDR <10^−7^, as reported by *clever*GO (Klus et al., 2015) as well as GeneMANIA server (https://genemania.org/) and includes SRSF 1, 5, 6 and 10, PCBP 1 and 2, HNRNP A0 and F, NOVA1, PPIG and TRA2A. *cat*RAPID *omics* predictions are in agreement with protein array experiments: from low- to high-affinity interactions, *cat*RAPID performances increase reaching AUCs of 0.80 (Figure 3C), indicating strong predictive power. Notably, while KHDRBS1 (not present in the protein array) is predicted to have poor binding propensity to CGG repeats, two of its RBP partners, CIRBP and PTBP2, rank in the top 1% of all fluorescence intensities, as predicted by *cat*RAPID (Cirillo et al., 2013), and DGCR8, which interacts with KHDRBS1 through DROSHA (Sellier et al., 2013), is found to interact (top 7% of all fluorescence intensities).

Out of 27 high-confidence candidates, 24 were predicted by *cat*GRANULE (Bolognesi et al., 2016) to form granules and among them the splicing regulator TRA2A showed the highest score (granule propensity = 2.15; Figure 3D; **Supplementary Figure 4D; Table S3**). In agreement with our predictions, eCLIP experiments indicate that the *FMR1* transcript ranks in the top 25% of strongest interactions with TRA2A (Van Nostrand et al., 2016a).

### TRA2A recruitment in FMR1 inclusions is driven by CGG hairpins in vivo

As splicing defects have been reported to occur in FXTAS disease (Botta-Orfila et al., 2016; Sellier et al., 2010), we decided to further investigate the recruitment of the splicing regulator TRA2A. B-lymphocytes are often used for initial investigations because of their easy accessibility from blood samples from patients. Expansions of CGG from 55 to 200 CGG repeats result in mRNA levels in B-lymphocytes that can exceed by 2–10 folds (Tassone et al., 2007). Therefore, B-lymphocytes are considered a good model to recapitulate some of the events occurring due to the permutation (i.e. higher expression levels of FMR1), and to explore new biomarkers. We measured RNA and protein levels of TRA2A in B-lymphocytes of a normal individual (41 CGG repeats; Coriell repository number NA20244A) and a FXTAS premutation carrier (90 CGG repeats; Coriell repository number GM06906B). RNA and protein levels of TRA2A were found significantly increased 2.9 and 1.4 times in the FXTAS premutation carrier compared to normal individual, which indicates that the TRA2A is significantly altered in disease (**Supplementary Figure 5**).

Yet, nuclear inclusions do not form in B-lymphocytes and we used the COS-7 cellular model to study *FMR1* inclusions (Sellier et al., 2010). We observed that transfection of a plasmid containing CGG expansions (triplet repeated 60 times) induce significant increase in RNA and protein levels of TRA2A after 48 hours (**Supplementary Figure 5**) (Sellier et al., 2010). By means of RNA FISH coupled to immunofluorescence (**STAR Methods**), we found that CGG expansions and endogenous TRA2A significantly co-localize in nuclear inclusions (45 out 50 screened cells showed unambiguous match). By contrast, TRA2A shows a diffuse nuclear pattern in cells that do not over-express CGG repeats (Figure 4A).

**Figure 4.**
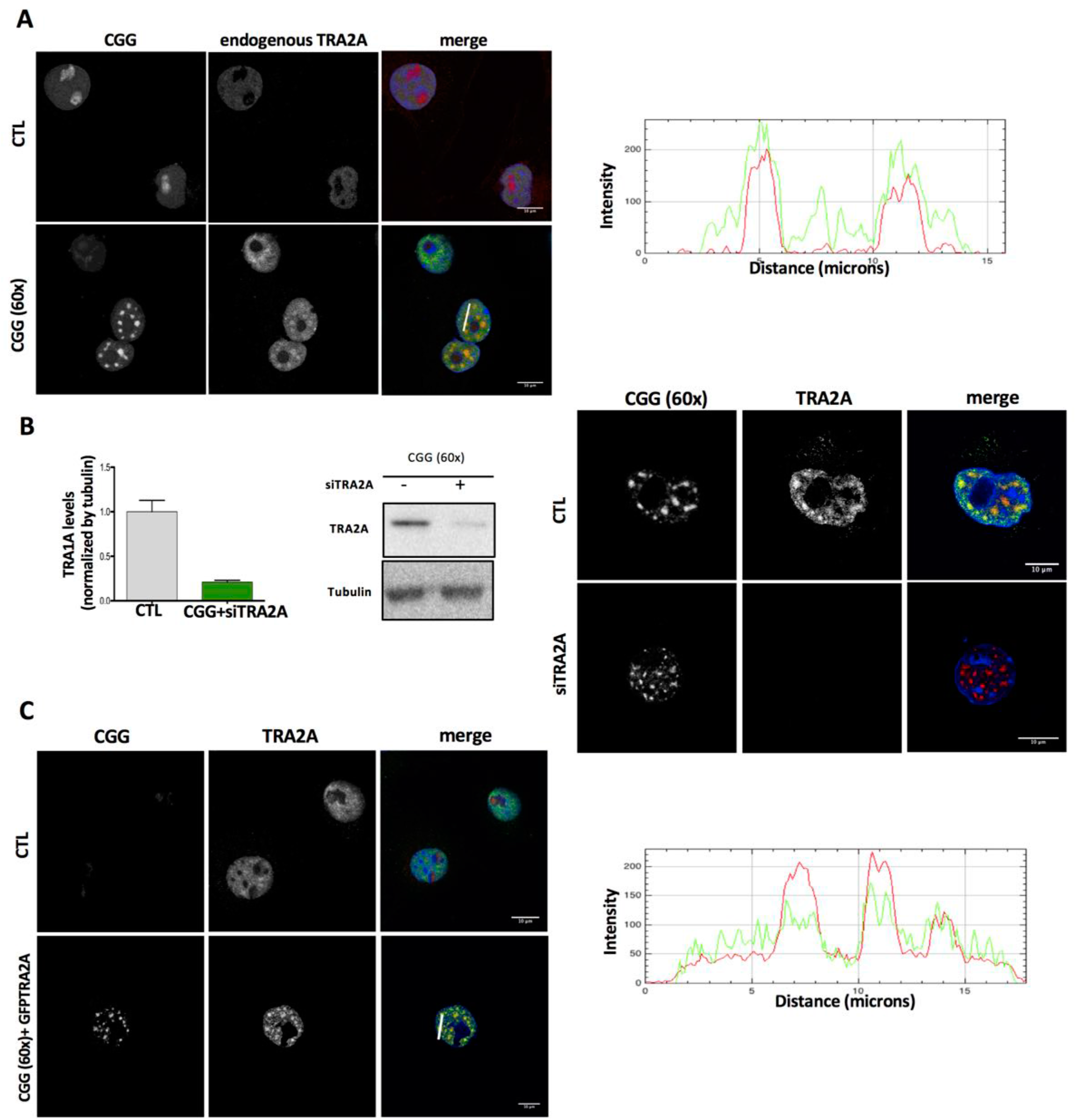
Endogenous TRA2A is recruited in nuclear RNA inclusions upon CGG over-expression. This specific recruitment is validated by experiments with TRA2A over-expression and TRA2A knockdown. **A)** COS-7 cells were transfected with either CGG(60X) or the empty vector as control. After 24h of transfection cells were immunostained with primary antiTRA2A antibody and secondary 488 and hybridized with a Cy3-GGC(8X) probe for RNA FISH. The graph represents the 488/Cy3 intensities co-localization in the section from the white line. **B)** After 24h of transfection cells were immunostained with antiTRA2A antibody and hybridized with a Cy3-GGC(8X) probe for RNA FISH; Relative TRA2A protein levels in COS-7 cells treated as in B. **C)** COS-7 cells were transfected with empty vector or CGG(60X) and GFP-TRA2A. After 48h, cells were hybridized with Cy3-GGC(8X) probe for RNA FISH. The graph represents the GFP/Cy3 intensities co-localization in the section from the white line.

Upon knockdown of TRA2A using siRNA (**STAR Methods**) we observed that the nuclear aggregates still form (Figures 4B and 4C), while over-expression of TRA2A attached to GFP (GFP-TRA2A) result in strong recruitment in CGG inclusions (Figure 4D; control GFP plasmid and GFP-TRA2A in absence of CGG repeats does not give a granular pattern).

To further characterize the recruitment of TRA2A in CGG repeats, we treated COS-7 cells with two different chemicals. By incubating COS-7 cells with 9-hydroxy-5,11-dimethyl-2-(2-(piperidin-1-yl)ethyl)-6H-pyrido[4,3-b]carbazol-2-ium (also named *1a*) that binds to CGG repeats preventing interactions with RBPs (Disney et al., 2012), TRA2A recruitment was blocked (Figure 5A). Using TmPyP4 to specifically unfold CGG repeats (Morris et al., 2012), we found that the aggregates are disrupted and TRA2A remains diffuse (Figure 5B).

**Figure 5.**
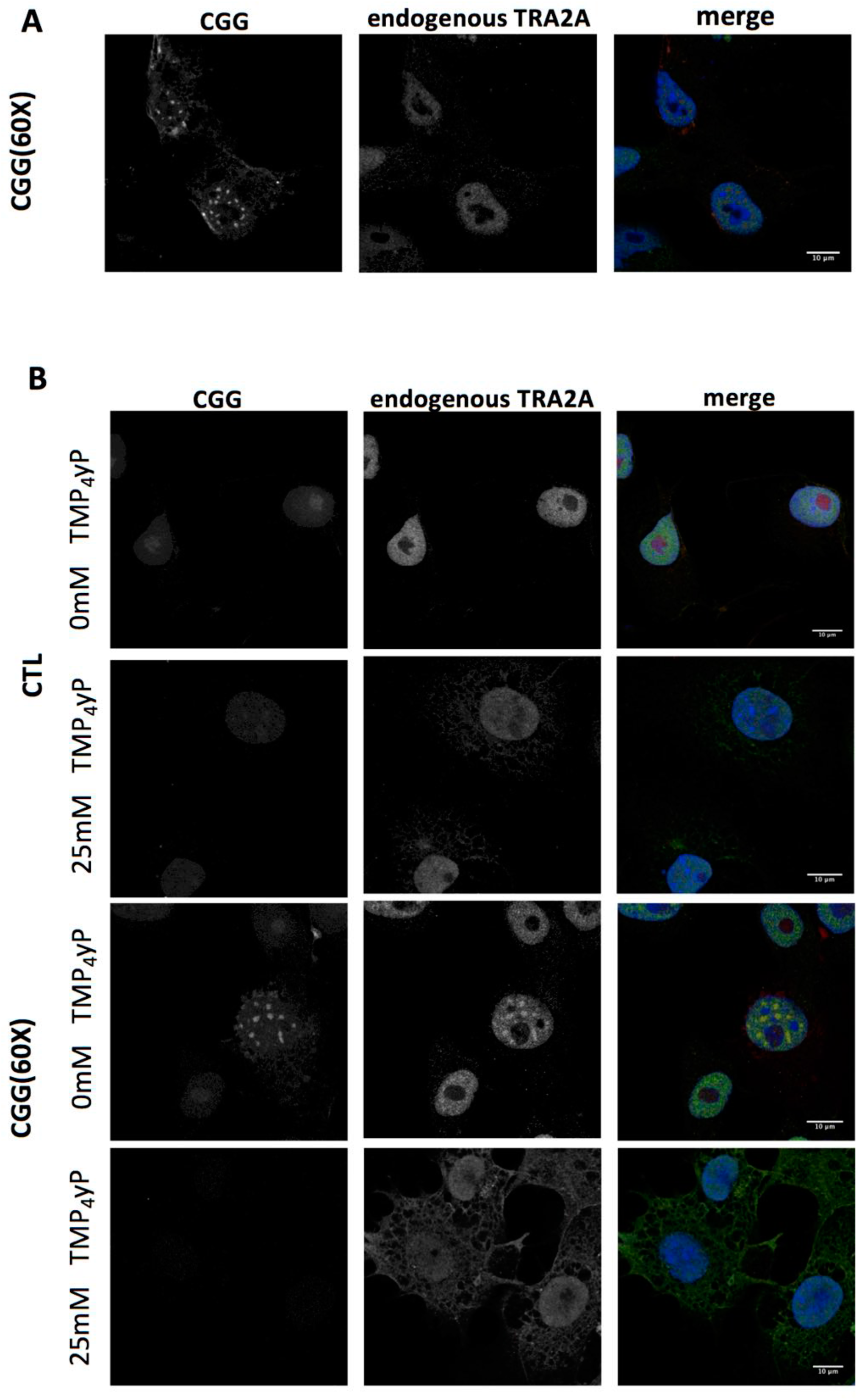
Disrupting CGG hairpins and dissolving RNA inclusions impair TRA2A sequestration. **A)** COS-7 cells were co-transfected with empty vector or CGG(60X), and after 24h of transfection cells were treated with 1a to block protein binding. **B)** COS-7 cells were treated similarly as in **A)** but with TmPyP4 molecule instead of 1a to disrupt CGG structure. In both cases, cells were immunostained with primary anti TRA2A antibody and hybridized with Cy3-GGC(8X) probe for RNA FISH.

Our experiments show that the aggregation of TRA2A is caused by CGG repeats and tightly depends on the hairping structure.

### TRA2A recruitment in RNA inclusions is independent of its partner TRA2B

Using RNA FISH coupled to immunofluorescence, we found that TRA2B, which interacts with TRA2A (Huttlin et al., 2015a) and binds to CGG repeats (Sellier et al., 2010), aggregates in COS-7 cells transfected with CGG repeats (x60; Figure 6A). Notably, endogenous TRA2B is recruited by CGG inclusions upon TRA2A knock-down (Figure 6B upper row**; Figure 6C;** the result is also observed when TRA2B is over-expressed; **Supplementary Figure 6A, 6B**). Similarly, endogenous TRA2A co-localizes with CGG repeats upon TRA2B knock-down (Figure 6B lower row; see also **Supplementary Figure 6C-E**). Thus, upon knock-down of one of the two proteins, the other one is still recruited by the over-expressed CGG repeats (Figure 6B and **Supplementary Figure 6F, 6G**). By contrast, in absence of CGG over-expression, neither TRA2A nor TRA2B localize within the inclusions (**Supplementary Figures 6A-E**).

**Figure 6.**
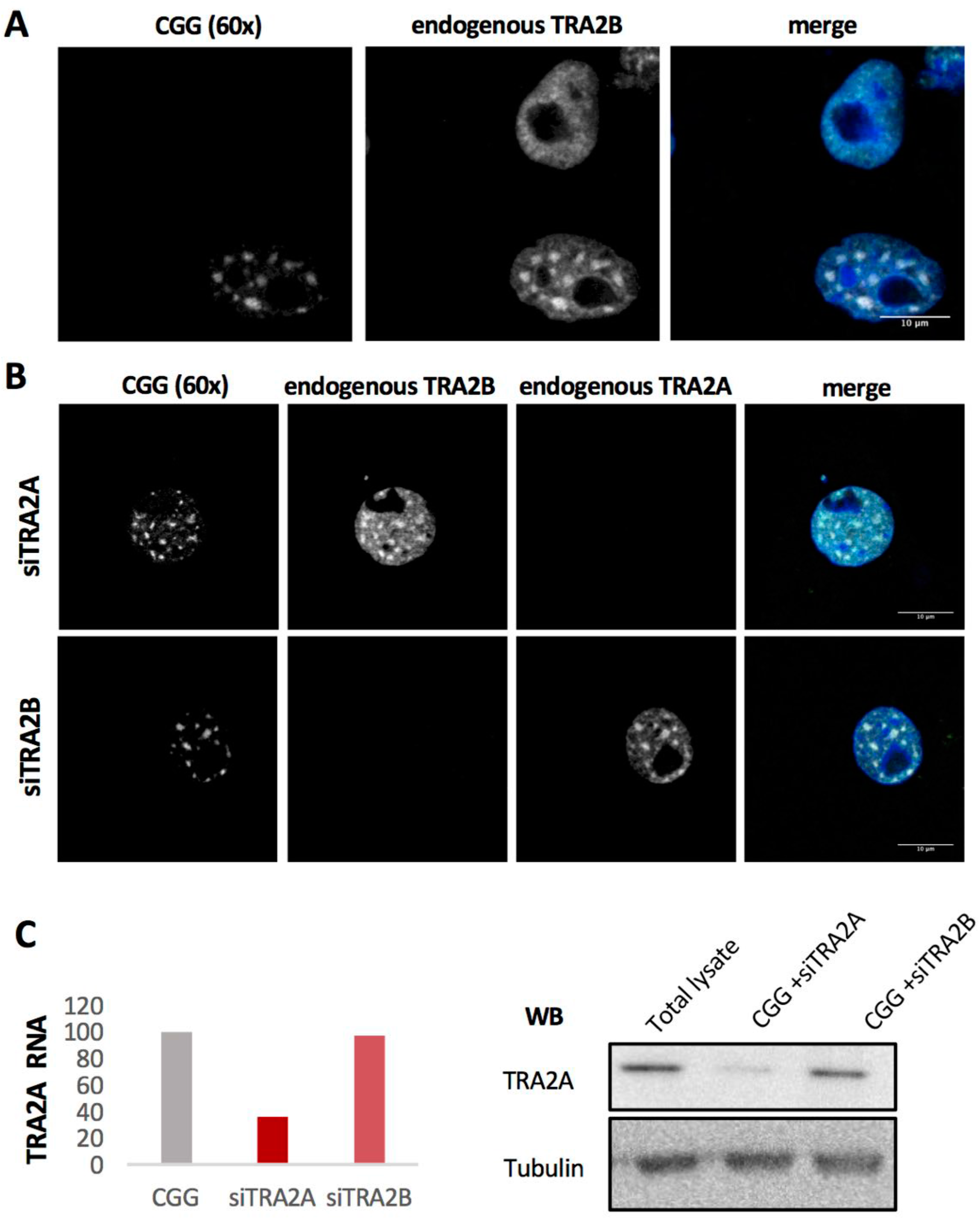
Endogenous TRA2B is recruited in CGG inclusions but TRA2A recruitment is independent from TRA2B. **A)** COS-7 cells were transfected with CGG(60X). After 24h of transfection cells were immunostained with antiTRA2B antibody and hybridized with a Cy3-GGC(8X) probe for RNA FISH. **B)** COS-7 cells were transfected with CGG(60X) and siTRA2A or siTRA2B. After 24h of transfection cells were immunostained with antiTRA2 antibodies and hybridized with a Cy3-GGC probe for RNA FISH. **C)** TRA2A protein levels in COS-7 cells treated as in **B**.

### Alterations in RNA splicing by FMR1 inclusions correlates with alterations in RNA splicing by TRA2A knockdown

When assessing TRA2A levels in response to CGG overexpression in COS-7 cells, we found that around 20-25% of TRA2A is recruited in condensates that are positive for CGG FISH signal (**STAR Methods**), which is fully compatible with a previous report on SOD1 accumulation in stress granules (Mateju et al., 2017). To study the functional implications of TRA2A recruitment in *FMR1* inclusions we analyzed changes in RNA splicing (**STAR Methods**).

Splicing alterations due to TRA2A sequestration were investigated through microarrays and RNA-seq experiments (both in triplicate experiments) to identify events (i) occurring upon CGG aggregates formation (CGG^+^ TRA2A^+^; 74 instances) and (ii) altered when TRA2A is knocked-down (CGG^+^ TRA2A^-^, 82 instances). With respect to events occurring in absence of CGG aggregates (CGG^-^ TRA2A^+^, i.e. physiological conditions), 59 exons are spliced in CGG^+^ TRA2A^+^ and not in CGG^+^ TRA2A^-^ (CGG^+^ TRA2A^+^ \ CGG^+^ TRA2A^-^) and thus depend on TRA2A sequestration (39 skipped and 20 included exons; q-value<0.10; Figure 7A; **Table S5**). Notably, 67 events occur exclusively in CGG^+^ TRA2A^-^ and can be ascribed to perturbations in the splicing factor network (Tan and Fraser, 2017), while 15 (i.e., 3+12) occur in both CGG^+^ TRA2A^+^ and CGG^+^ TRA2A^-^ (Figure 7B) and are therefore TRA2A independent (Figure 7B).

**Figure 7.**
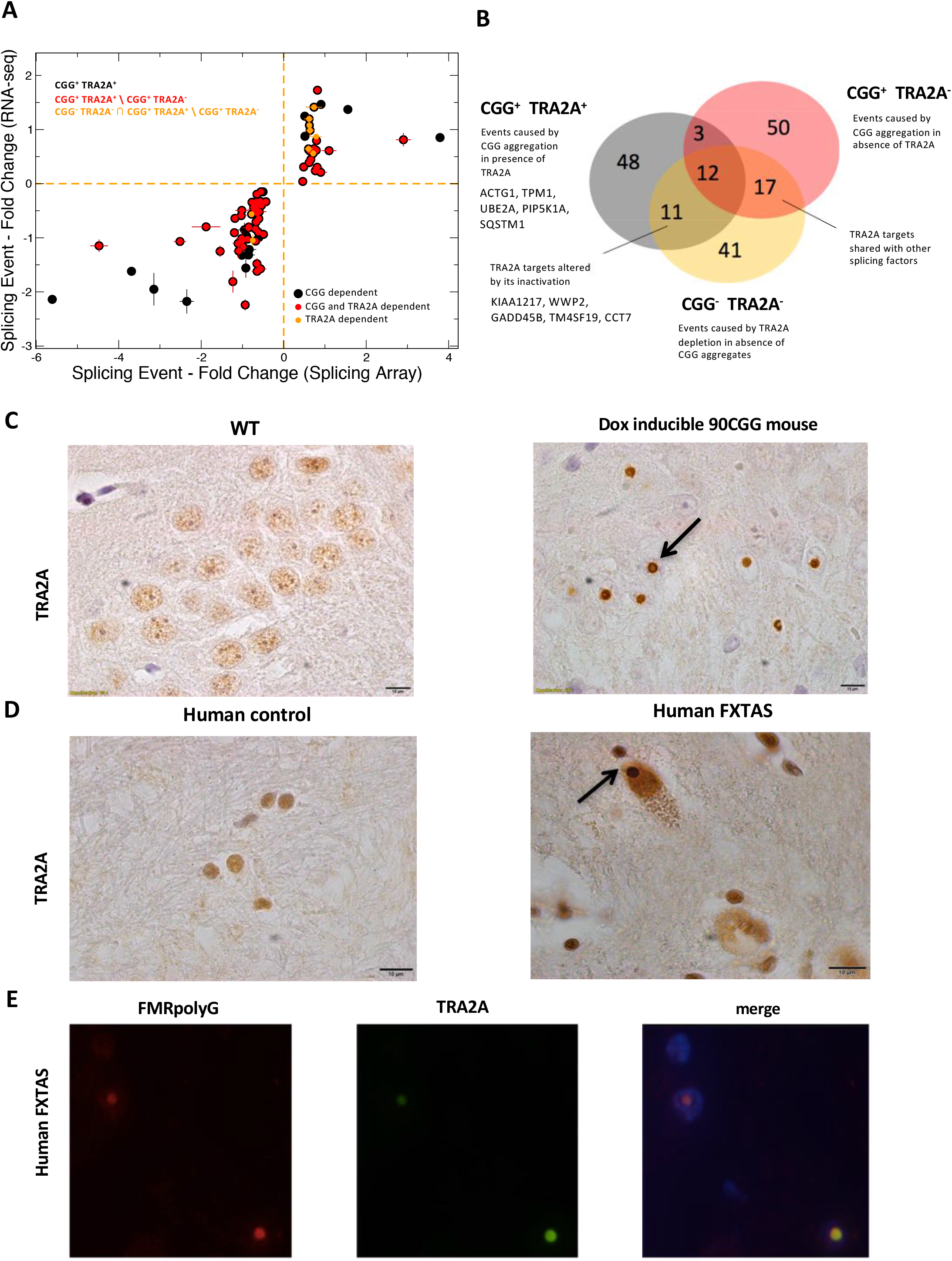
TRA2A recruitment in *FMR1* inclusions affects splicing. **A)** With respect to events occurring in absence of CGG aggregates (CGG^-^ TRA2A^+^), 59 exons are spliced in CGG^+^ TRA2A^+^ and not in CGG^+^ TRA2A^-^ (i.e., CGG^+^ TRA2A^+^ \ CGG^+^ TRA2A^-^; red points). We investigated cases caused by TRA2A depletion in absence of CGG aggregates (CGG^-^ TRA2A^-^) and identified 11 events that are present in CGG^+^ TRA2A^+^ and not in CGG^+^ TRA2A^-^ (i.e.; (CGG^-^ TRA2A^-^ **∩** CGG^+^ TRA2A^+^ \ CGG^+^ TRA2A^-^; orange points; **Table S5; STAR Methods**). **B)** Muscle proteins, including PIP5K1A, TPM1 and autophagy-related SQSTM1 are subjected to exon skipping upon TRA2A recruitments in RNP condensates (**Table S5**; Figure 7B). Events occuring when TRA2A is depleted are linked to neuro-pathogenesis (GADD45B and CCT7) and skeletal development (KIAA1217 and TM4SF19). **C)** TRA2A immunohistochemistry in wild type (WT) and premutated mouse model (counterstaining is done with haematoxylin). **D)** TRA2A immunohistochemistry in human hippocampus from control and FXTAS (counterstained with haematoxylin; the arrows points to the inclusion). **E)** Double immunofluorescence of TRA2A as well as FMRpolyG peptides in human FXTAS (**STAR Methods**).

To better understand TRA2A-dependent effects, we studied events caused by TRA2A depletion in absence of CGG aggregates (CGG^-^ TRA2A^-^; Figure 7A): 11 out of 59 CGG^+^ TRA2A^+^ cases are present in CGG^-^ TRA2A^-^ but not in CGG^+^ TRA2A^-^ (CGG^-^ TRA2A^-^ **∩** CGG^+^ TRA2A^+^ \ CGG^+^ TRA2A^-^), which is statistically highly significant. Indeed, by shuffling the splicing events reported in one of the two experiments (i.e., randomizing the association between exons and q-values), we found that the intersection CGG^-^ TRA2A^-^ **∩** CGG^+^ TRA2A^+^ \ CGG^+^ TRA2A^-^, never contains 11 events (we found 1 event in 288 out 10000 randomizations and 2 events in 3 out 10000 randomizations, but never >3 events; p-value << 10^−4^; Figure 7B). Notably, 17 cases occur in CGG^+^ TRA2A^-^ and CGG^-^ TRA2A^-^ but not in CGG^+^ TRA2A^+^, which is expected since splicing factors work together on common targets (Tan and Fraser, 2017).

Using the *clever*GO algorithm (Klus et al., 2015) we found that the largest GO cluster of affected genes includes RBPs (18 genes; **Table S5**; ‘RNA-binding’; fold enrichment of 24; p-value < 10^−8^; calculated with Bonferroni correction; examples: HNRNPL, CIRBP and DDX24) and, more specifically, spliceosome components (‘mRNA splicing via spliceosome’; fold enrichment of 5; p-value < 10^−3^, examples: HNRNP A2/B1 and SRSF 10) or genes related to alternative splicing activity (‘regulation of splicing’; fold enrichment of 6; p-value < 10^−3^, examples: RBM5 and THOC1).

Intriguingly, genes associated with mental retardation, such as UBE2A (Budny et al., 2010), ACTB (Procaccio et al., 2006) and ACTG1 (Rivière et al., 2012), have splicing patterns affected by TRA2A sequestration. Similarly, muscle related proteins, including PIP5K1A (Chen et al., 2018), TPM1 (Erdmann et al., 2003) and genes linked to intellectual disabilities such as DOCK3 (de Silva et al., 2003) and craniofacial development, such as WWP2 (Zou et al., 2011), are subjected to exon skipping upon TRA2A recruitments in RNP condensates (**Table S5**; Figure 7B). Out of 59 splicing events occurring in CGG^+^ TRA2A^+^ and CGG^+^ TRA2A^-^ conditions, 23 (including ACTG1, TMP1 and WWP2) involve transcripts that physically bind to FMRP protein, as also detected in CLIP experiments (available from http://starbase.sysu.edu.cn/), which unveils an important link (significance: p-value < 10^−4^; Fisher’s exact test) to Fragile X Syndrome (Maurin et al., 2014).

In the 11 CGG^+^ TRA2A^+^ cases present in CGG^-^ TRA2A^-^ but not in CGG^+^ TRA2A^-^ (Figure 7B) there is GADD45B linked to synaptic plasticity (Ma et al., 2009), as well as g KIAA1217 (Semba et al., 2006) and TM4SF19 (de la Rica et al., 2013) associated with skeletal development pathways. We also found the molecular chaperone CCT that is known to restrict neuro-pathogenic protein aggregation via autophagy (Pavel et al., 2016).

### TRA2A is present in murine and human FXTAS brain inclusions

We tested if TRA2A co-aggregates with *FMR1* inclusions in a mouse model in which the 5’UTR (containing 90 CGG repeats) was expressed under the control of doxycycline (Hukema et al., 2015). Immunohistochemistry experiments with sections of paraffin-embedded neurons and astrocytes indicated that TRA2A protein is present in the inclusions (Figure 7C; STAR Methods).

Importantly, repeat associated non-AUG (RAN) translation has been shown to occur in *FMR1* 5’UTR, resulting in the production of FMRpolyG and FMRpolyA peptides (Todd et al., 2013). The main RAN translation product, FMRpolyG, co-localizes with ubiquitin in intranuclear inclusions (Sellier et al., 2017).

In agreement with our murine model, we found positive staining for TRA2A in nuclear inclusions from two FXTAS *post mortem* human brain donors (Figure 7D), and remarkably, we observed co-localization with FMRpolyG (Figure 7E). We observe that FMRpolyG reaches its highest abundance in hippocampus co-aggregating in 20% total *nuclei* (Glineburg et al., 2018). In the same tissue, TRA2A co-localizes with inclusions in 2-3% of total *nuclei*, thus indicating strong sequestration (**Supplementary Figure 7**). Interestingly, TRA2A positive cells aggregate in groups that are in close proximity, which provides precious information on the biochemical behavior of aggregates as well as spreading nature of the disease across brain districts (**Supplementary Figure 7**).

Thus, TRA2A sequestration by CGG repeats is not only observed in cell lines, but also in FXTAS animal models and human *post mortem* brain samples.

## Discussion

Previous evidence indicates that proteins are the main cohesive elements within RNP granules (Banani et al., 2017). Yet, specific RNAs can act as structural scaffolds to assemble proteins in RNP condensates, as recently reported in literature (Langdon et al., 2018) and RNA-RNA interactions play an important role in the formation of RNP assemblies (Treeck and Parker, 2018). Our analysis of PRI networks reveals that scaffolding RNAs have large number of RBP contacts, increased length and high structural content. In agreement with our computational analysis, two works published at the time of writing indicate that UTRs length and structural content (Khong et al., 2017; Maharana et al., 2018) are important properties of RNAs aggregating in RNP condensates. Moreover, nucleotide repeats (Jain and Vale, 2017), changes in RNA levels (Tartaglia and Vendruscolo, 2009) and RNA binding abilities (Zhang et al., 2015) are known factors modulating phase transitions in the cell.

Our PRI networks were retrieved from eCLIP experiments (Van Nostrand et al., 2016a) that have been performed in conditions different from those promoting formation of physiological stress granules. Similarly, PPI networks were taken from the BioPlex database that includes highly curated multi-source experiments (Huttlin et al., 2015a). Yet, our underlying hypothesis is that PPI and PRI are governed by physico-chemical forces that are in place regardless of the environmental conditions and we assume that the ability of proteins and RNAs to assemble is impaired when the molecules are poorly expressed or chemically modified. Indeed, to control for the contribution of RNA abundance, we used it to normalize the number of CLIP reads in our calculations. Supporting our assumptions, the RNA list retrieved from the networks analysis shows a very significant overlap with a recently published atlas of transcripts enriched in SGs (Khong et al., 2017). We note that our analysis would be more accurate if the protein and RNA interactions networks were known for the different biological condensates.

Combining computational approaches with large-scale *in vitro* experiments we unveiled the scaffolding ability of *FMR1* 5’ UTR, recovering previously known partners relevant in FXTAS such as SRSF1, 5, 6, KHDRBS3 and MBNL1 (Sellier et al., 2010) and identifying additional interactions involved in alternative splicing, such as PCBP 1 and 2, HNRNP A0 and F, NOVA1, PPIG and TRA2A. At the time of writing, TRA2A has been reported to be a component of ALS granules (Markmiller et al., 2018). Yet, TRA2A does not appear in TAU inclusions (Maziuk et al., 2018), which indicates that its sequestration occurs only in specific neurodegenerative diseases.

To prove the implication of TRA2A sequestration in FXTAS pathogenesis, and overcome the technical limitation of our cellular model in which the non-AUG codon downstream the 5’UTR of *FMR1* is lacking, we tested TRA2A colocalization with FMRpolyG in patients’ brain samples. Our experiments showed that the two proteins colocalize, providing additional information on TRA2A involvement in FXTAS disease. This result indicate that CGG could interact with both TRA2A and FMRpolyG and supports our previous work indicating that interactions between proteins and cognate RNAs are frequent in aggregation-prone genes (Cirillo et al., 2013).

Through splicing microarrays and RNA-seq analysis we found that TRA2A sequestration induces changes in splicing of genes associated with mental retardation, including ACTB (Procaccio et al., 2006) and ACTG1 (Rivière et al., 2012), intellectual disabilities, such as DOCK3 (de Silva et al., 2003), and craniofacial development, such as WWP2 (Zou et al., 2011), which are relevant in the context of Fragile X Syndrome (Maurin et al., 2014). Thus, the identification of TRA2A opens the avenue for new therapeutic intervention to correct the splicing defects of deregulated transcripts or to restore the functional role by CRISPR/Cas technology.

In the future, it will be very important to analyze genome-wide data in different bio-specimens from patients to see the expression of differently spliced variants of TRA2A targets. Nevertheless, since *FXTAS* is a rare disease with low penetrance, the amount of samples from patients is very limited. Therefore, more work should be done in this direction to promote biomarker discovery in patients and ultimately promote personalized treatment. Yet, our theoretical framework is also applicable to other diseases in which RNAs promote formation of phase-separated condensates that could be used by the pathologist to identify the proteins that are specifically sequestered.

## ACKNOWLEDGEMENTS

The authors are deeply in debt to brain donors and their relatives. We acknowledge specimen donations from Dr Nicolas Charlet-Berguerand (pcDNA3 CGG 60X), Dr David Elliott and Dr Caroline Dalgliesh (plasmid GFP TRA2A), Dr Eulàlia Martí (plasmids CGG 21X and 79X and isolated fibroblasts from CAG carriers), Dr Matthew D. Disney (molecule *1a*). We thank Martin Vabulas for the “schemino” of Figure 1A and all members of Tartaglia’s lab, especially Dr. Elias Bechara for the interpretation of splicing experiments. We are grateful to Dr Lara Nonell Mazelón and Dr Magdalena Arnal Segura for RNAseq and splicing arrays analysis, Dr Fatima Gebauer, Dr Natàlia Sánchez de Groot, Dr Davide Cirillo and Dr Domenica Marchese for stimulating discussions.

The research leading to these results has been supported by European Research Council (RIBOMYLOME_309545), Spanish Ministry of Economy and Competitiveness (BFU2014-55054-P and BFU2017-86970-P) and “Fundació La Marató de TV3” (PI043296). We acknowledge support of the Spanish Ministry of Economy and Competitiveness, ‘Centro de Excelencia Severo Ochoa 2013-2017’. We acknowledge the support of the CERCA Programme / Generalitat de Catalunya. Support of Spanish Ministry for Science and Competitiveness (MINECO) to the EMBL partnership.

## Author contributions

GGT and TBO conceived the study together with the help of BB, FCS performed the calculations, TBO supervised MGB and performed all the experiments as well as analyzed samples from RH, LWS and EG. TBO, NLG, GGT and BL analyzed the data. TBO, FCS, BB and GGT wrote the manuscript.

